# Spike-timing-dependent plasticity rewards synchrony rather than causality

**DOI:** 10.1101/863365

**Authors:** Margarita Anisimova, Bas van Bommel, Marina Mikhaylova, J. Simon Wiegert, Thomas G. Oertner, Christine E. Gee

**Author notes:** Correspondence: Christine Gee, Institute for Synaptic Physiology, Center for Molecular Neurobiology Hamburg, Falkenried 94, 20251 Hamburg, Germany. These authors contributed equally to this work (Thomas G. Oertner, Christine E. Gee).

## Abstract

Spike-timing-dependent plasticity (STDP) is a candidate mechanism for information storage in the brain, but the whole-cell recordings required for the experimental induction of STDP are typically limited to one hour. This mismatch of time scales is a long-standing weakness in synaptic theories of memory. Here we use spectrally separated optogenetic stimulation to fire precisely timed action potentials (spikes) in CA3 and CA1 pyramidal cells. Twenty minutes after optogenetic induction of STDP (oSTDP), we observed timing-dependent depression (tLTD) and timing-dependent potentiation (tLTP), depending on the sequence of spiking. As oSTDP does not require electrodes, we could also assess the strength of these paired connections three days later. At this late time point, late tLTP was observed for both causal (CA3 before CA1) and anti-causal (CA1 before CA3) timing, but not for asynchronous activity patterns (Δt = 50 ms). Blocking activity after induction of oSTDP prevented stable potentiation. Our results confirm that neurons wire together if they fire together, but suggest that synaptic depression after anti-causal activation (tLTD) is a transient phenomenon.

**Highlights:**

- Optogenetic induction of spike-timing-dependent plasticity at Schaffer collateral synapses
- Causal pairing induces potentiation whereas anti-causal pairing induces depression during patch-clamp recordings.
- Three days after optogenetic induction, the consequence of STDP is potentiation (tLTP) irrespective of spiking order.
- Late tLTP requires ongoing activity in the days following oSTDP.

**Graphical Abstract:** 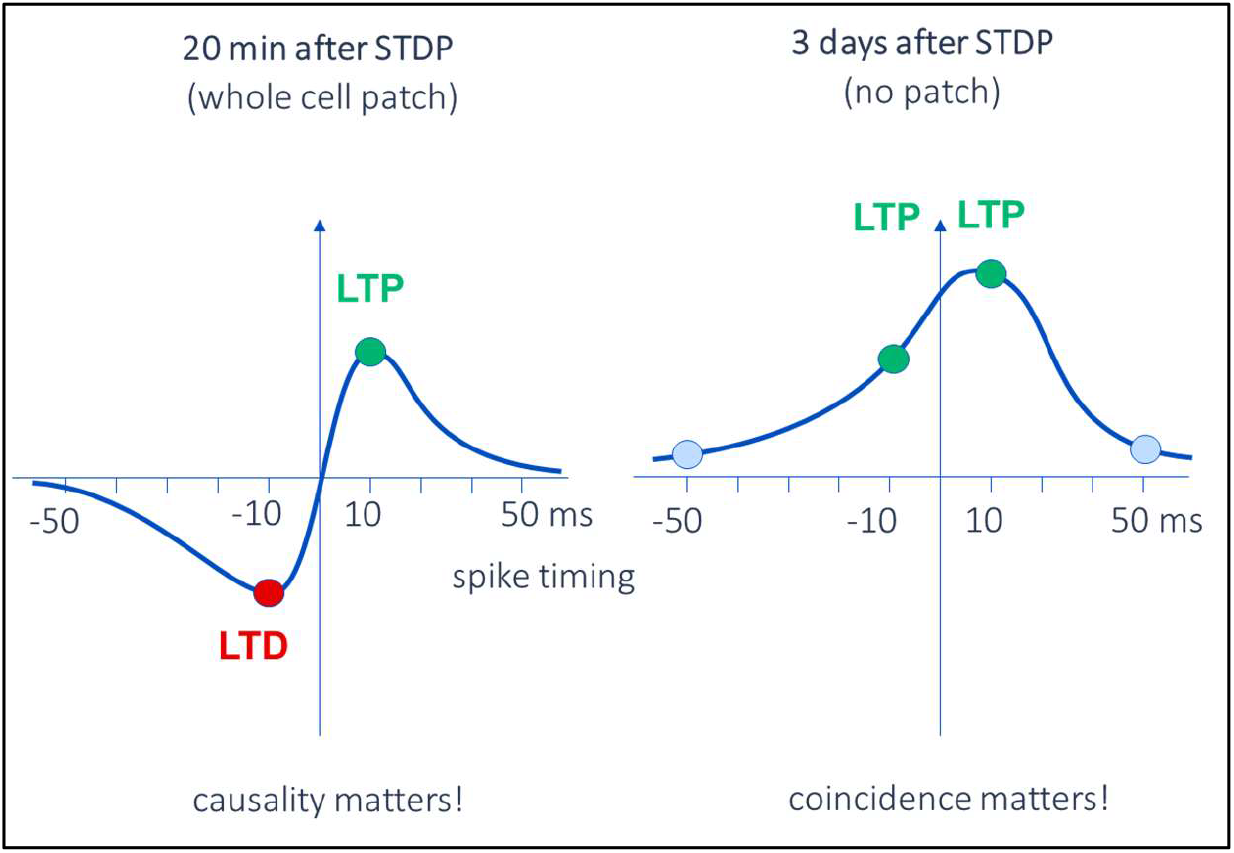

## Introduction

Synapses change their strength in response to synchronized activity in pre- and postsynaptic neurons. This fascinating property, commonly referred to as spike-timing-dependent plasticity (STDP), has inspired numerous computational models of memory formation ^1–4^. Indeed, STDP-like learning rules are now widely used for unsupervised learning, have been implemented on neuromorphic hardware and in robotics ^5–7^. Canonical STDP bidirectionally changes synaptic strength depending on the order of pre- and postsynaptic activity. Timing-dependent long-term potentiation (tLTP) occurs when excitatory synaptic potentials repeatedly precede action potentials, i.e. causal or Hebbian pairing. In contrast, timing-dependent long-term depression (tLTD) occurs when this order is reversed, i.e. anti-causal pairing ^8–10^. Depending on the synapse in question, the exact spiking pattern and the presence or absence of neuromodulators, different STDP rules have been described ^11,12^. Typically, STDP experiments follow changes in synaptic strength for 20-60 minutes after induction. This time limitation arises from the invasive nature of intracellular recordings, which are necessary to control the precise timing of postsynaptic spikes. Memories last much longer, and it has always been conceptually difficult to reconcile the millisecond precision required for STDP mediated synaptic strengthening with memories of associated events, which form regardless of exact order of presentation. Our aim was therefore to extend the post-STDP observation time from hours to several days and examine the late effects on synaptic strength.

Channelrhodopsins are light-activated ion channels, and since the discovery of channelrhodopsin-2 ^13^ have been widely used to stimulate neurons *in vitro* and *in vivo* ^e.g. 14–16^. In recent years, channelrhodopsin variants with different kinetics and spectral sensitivities have been discovered or engineered ^e.g. 17,18^. Here, we present a two-color method to evoke precisely timed action potentials in defined subsets of CA3 and CA1 pyramidal neurons for optogenetic induction of STDP (oSTDP). By combining the very light-sensitive opsin CheRiff and less sensitive ChrimsonR, we could independently drive precisely timed action potentials in CA3 and CA1 pyramidal neurons using dim 405 nm and bright 625 nm light pulses, respectively. As expected, we found that during short-term patch-clamp recordings (20 minutes after induction), oSTDP resulted in tLTP after causal (single presynaptic spike before postsynaptic burst) pairing and tLTD after anti-causal (postsynaptic burst before single presynaptic spike) pairing, a classical asymmetric plasticity window. However, 3 days after oSTDP, the window became fairly symmetric, comprised of tLTP at short (±10 ms) delays and no change with longer (±50 ms) delays (i.e., phase-shifted activity). Late tLTP depended on pairing frequency (at 5 Hz but not at 0.1 Hz) and activation of NMDA-type glutamate receptors. We also observed that the appearance of late tLTP after anti-causal pairing requires ongoing spontaneous activity in the network. Our findings reveal that short-term recordings of STDP cannot simply be extrapolated to predict synaptic strength over behaviorally relevant time scales. Especially the frequently reported tLTD might not be ‘long-term’ after all.

## Results

### Independent optogenetic control of pre- and postsynaptic action potentials

To independently spike pre- and postsynaptic neurons, we used the red light sensitive ChrimsonR ^17^ and the blue/violet light-sensitive CheRiff ^18^. ChrimsonR-tdTomato was expressed in a small subset of CA3 pyramidal neurons by local AAV injection and several CA1 neurons were electroporated with DNA encoding CheRiff-eGFP (Fig. 1A). Based on calibration experiments (Fig. 1B, C), we selected 405 nm (1 mW mm^-2^) to activate CheRiff-CA1 neurons without activating ChrimsonR-CA3 neurons, and 625 nm (8 mW mm^-2^) to activate ChrimsonR-CA3 neurons. These intensities reliably and selectively evoked single action potentials/spikes in transduced CA3 or CA1 pyramidal neurons using 1-2 ms flashes from 7 to 10 days after transduction (e.g. Fig. 1D). As CheRiff-CA1 neurons were completely insensitive to wavelengths longer than 565 nm, we co-expressed the red fluorescent protein mKate2 to visualize them without spiking the postsynaptic neurons.

**Figure 1.**
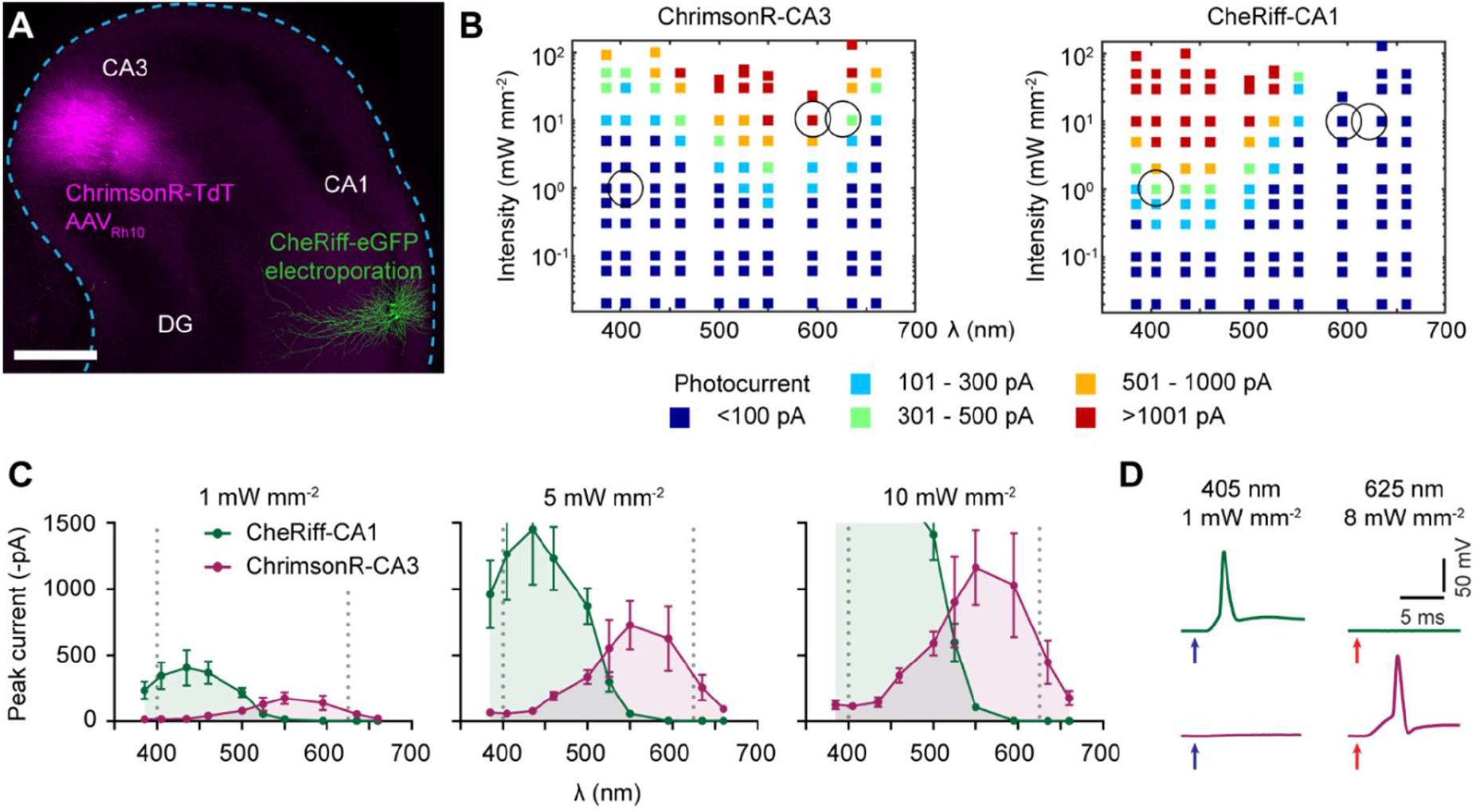
Selecting wavelength and intensity for independent spiking of CheRiff-CA1 and ChrimsonR-CA3 neurons. **A**, Confocal image (maximum intensity projection) of an organotypic hippocampal slice culture (DIV 21) taken 7 days after transducing ∼30 CA3 neurons with AAVRh10-syn-ChrimsonR-tdTomato (magenta) and electroporating 4 CA1 neurons with pAAV-syn-CheRiff-eGFP (green). Scale bar 500 µm. **B**, Color-coded photocurrent amplitude in response to 1 ms light flashes of varying intensity and wavelength of ChrimsonR-CA3 (left) and CheRiff-CA1 neurons (right). Circles indicate parameters used in plasticity experiments. n = 4 recordings per point, from 11 ChrimsonR-CA3 and 7 CheRiff-CA1 neurons. **C**, Photocurrent amplitude vs wavelength from the data in B at intensities of 1, 5 and 10 mW mm^-2^. Regions of overlap will depolarize both CheRiff- and ChrimsonR-expressing neurons. At 10 mW mm^-2^, the maximum current recorded from CheRiff-CA1 neurons was -2100 ± 500 pA at 435 nm (mean ± SEM). Dotted lines indicate the optimal wavelengths to independently activate ChrimsonR-CA3 and CheRiff-CA1 neurons. Mean ± SEM, n = 4 measurements per point. **D**, Example membrane responses of a CheRiff-CA1 (upper traces) and a ChrimsonR-CA3 (lower traces) neuron to 2 ms light flashes at 405 nm (1 mW mm^-2^, blue arrows) and 625 nm (8 mW mm^-2^, red arrows). Note the CheRiff-CA1 neuron fires a single action potential and the ChrimsonR-CA3 neuron only slightly depolarizes in response to 405 nm light, whereas the ChrimsonR-CA3 neuron fires one action potential in response to 625 nm and there is no response in the CheRiff-CA1 neuron.

### Optogenetic induction of spike-timing-dependent plasticity

As proof of principle, we induced STDP all-optically while patched on the postsynaptic neuron (Fig. 2A, B). We recorded excitatory postsynaptic currents (EPSCs) in CheRiff- or non-transfected CA1 (NT-CA1) neurons while optogenetically stimulating the ChrimsonR-CA3 neurons through the condenser with light flashes at 20 s intervals (594 nm). After collecting baseline responses for 5 min, we switched from voltage-to current-clamp mode. Single presynaptic ChrimsonR-CA3 spikes (300 flashes at 5 Hz) evoked EPSPs that preceded or followed three-spike bursts by about 10 ms in the transfected CheRiff-CA1 postsynaptic neurons (Tr, 3 flashes at 50 Hz, Fig. 2C). Causal pairing (pre-before post-) induced tLTP (n = 12 experiments; p = 0.005, paired t-test, Fig, 2D, F, for details of statistical analyses see Suppl. Table 1) whereas anti-causal pairing (post-before pre-) induced tLTD (n = 11 experiments, p = 0.02, paired t-test, Fig. 2E, F). The normalized changes in synaptic strength after causal and anti-causal pairing were significantly different (Fig. 2F, p = 0.0003, ANOVA-Sidak). Importantly, there was no change in synapses onto NT-CA1 neurons (n = 6 experiments, p = 0.9, paired t-test, Fig. 2F), indicating that oSTDP was specific to synapses between co-activated CA3 and CA1 neurons and no plasticity was induced by the 5 Hz presynaptic stimulation alone. Thus, using light to induce spike-burst STDP produces the typical asymmetric tLTD-tLTP window at Schaffer collateral synapses ^8,9,19–23^.

**Figure 2.**
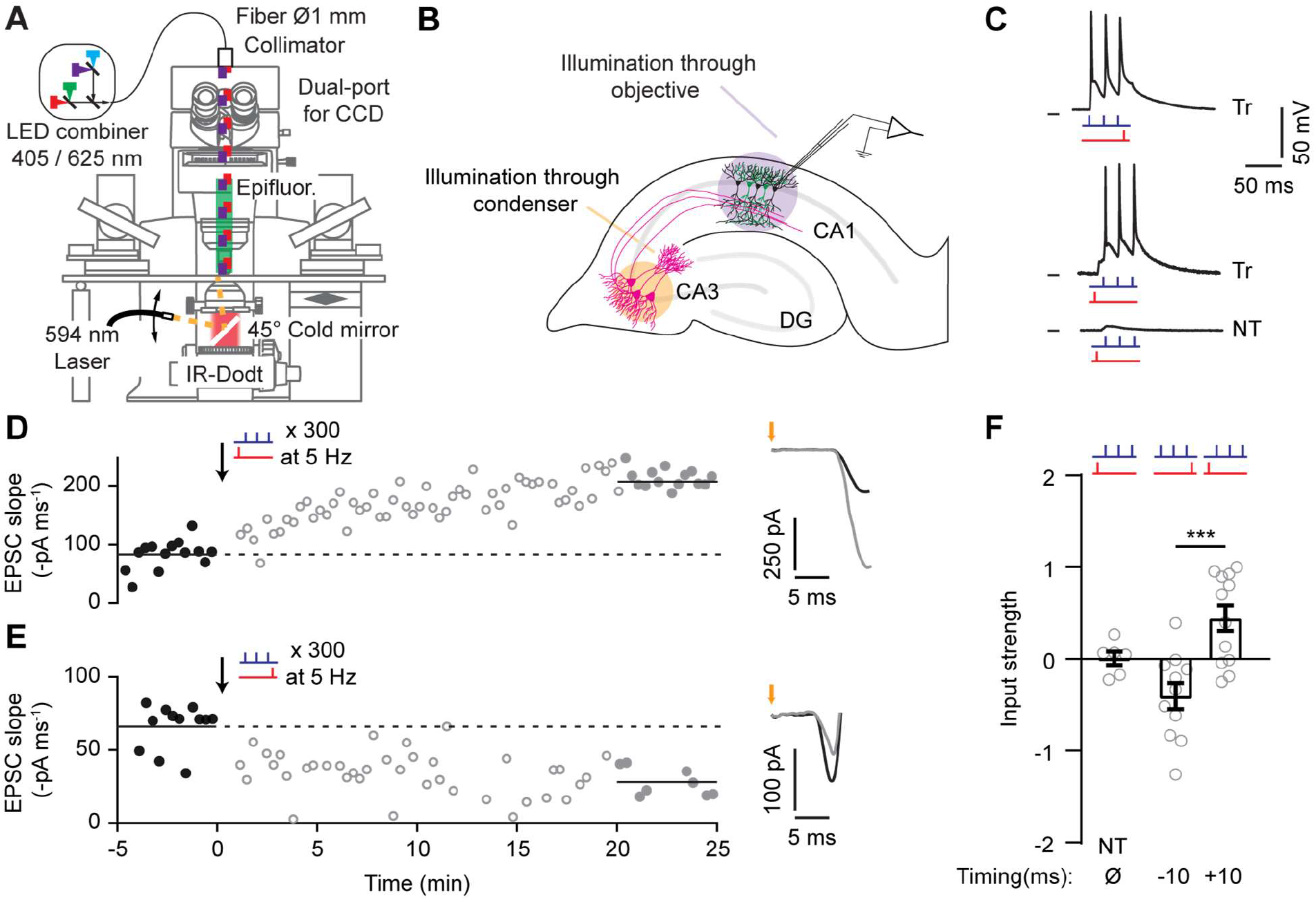
Optogenetic induction of spike-timing-dependent plasticity (oSTDP) produces causality-dependent tLTP and tLTD. **A**, Diagram of the electrophysiological recording setup for on-axis CA1 stimulation through the objective and off-axis CA3 stimulation through the condenser. **B**, Experimental configuration with patch-electrode in CA1. **C**, Current-clamp recordings from CA1 neurons during oSTDP induction. Top, a CheRiff-transfected (Tr) CA1 neuron during anti-causal pairing (−10 ms: 3 violet (405 nm) flashes at 50 Hz and 1 red (625 nm) flash 8 ms after, repeated 300x at 5 Hz). Middle, a CheRiff-transfected CA1 neuron during causal pairing (+10 ms: 1 red flash and 3 violet flashes at 50 Hz 12 ms after). Bottom, a non-transfected (NT) CA1 neuron during causal pairing. Black ticks at left indicate -70 mV. **D**, Left, example causal pairing experiment from one CheRiff-CA1 neuron. Excitatory postsynaptic currents (EPSC) were evoked by light stimulation of ChrimsonR-CA3 neurons before (black points) and after (grey points) causal pairing at t = 0 (black arrow). The filled points were significantly different, p < 0.0001, Kolmogorov-Smirnov. Right, averaged EPSCs from the filled points, orange arrow indicates stimulation of ChrimsonR-CA3 neurons. **E**, As in panel **D**, but after anti-causal stimulation (black arrow). The filled gray points were significantly different to the baseline, p = 0.0003, Kolmogorov-Smirnov. **F**, Normalized change in EPSC slope 20-25 min after oSTDP induction as in panels **D** (+10, n = 12) and **E** (−10, n = 11). NT, non-transfected neurons from slices subjected to causal pairing stimulation (n = 6). *** p = 0.0003, ANOVA-Sidak.

### Input strength is potentiated three days after no-patch oSTDP

To induce STDP inside the incubator, we constructed illumination towers containing independently controlled, collimated red (625 nm) and violet (405 nm) high-power LEDs to stimulate ChrimsonR-CA3 and CheRiff-CA1 neurons with light pulses of defined intensity (Fig. 1). After causal pairing, we assessed expression of the immediate early gene cFos, which is upregulated in burst-spiking neurons ^24^ and in neurons that have undergone LTP ^for review see 25^. Eighty-three percent of CheRiff-CA1 neurons expressed cFos, indicating they were spiking in bursts during causal pairing as expected (7 slices: 30/36 neurons; e.g. Fig. 3A). Whether non-transfected (NT) CA1 neurons also expressed cFos depended critically on the number of ChrimsonR-CA3 neurons in the slice. In 5 slices with 36-53 ChrimsonR-CA3 neurons, no NT-CA1 neurons expressed cFos (Fig. 3A). In 2 slices with 60-61 ChrimsonR-CA3 neurons, some NT-CA1 neurons showed cFos staining, and in 2 slices with 96 or more ChrimsonR-CA3 neurons, cFos staining was evident throughout the entire CA1 (Fig. 3B, C). Based on these observations, we adjusted the amount of ChrimsonR virus injected to transduce ∼30 CA3 neurons per slice culture (e.g. Fig 1A) because to induce STDP at a select subset of synapses, it is important to avoid spike bursts in non-transfected neurons.

**Figure 3.**
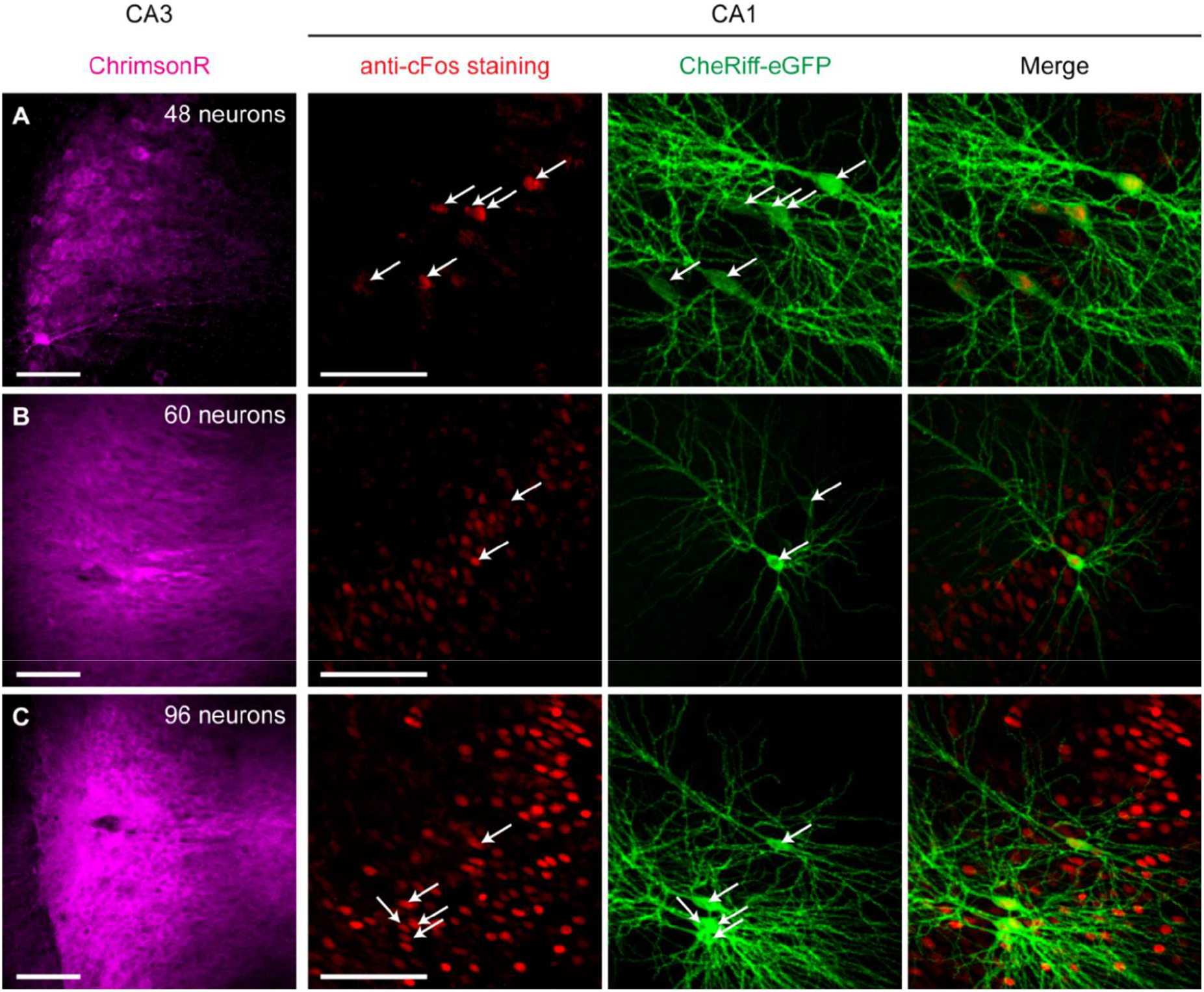
CA1 cFos expression after causal pairing depends on the number of ChrimsonR-CA3 neurons. Confocal images (average intensity projection) of CA3 and CA1 areas of three slice cultures expressing ChrimsonR-tdT in CA3 (magenta) and CheRiff-eGFP in CA1 neurons (green, white arrows indicate nuclei). In red is cFos immunofluorescence. Slices were fixed one hour after causal pairing (see Fig 2D). **A**, A slice with fewer than 50 ChrimsonR-CA3 neurons. Note that cFos immunoreactivity is restricted to the CheRiff-CA1 neurons (5 of 6 are cFos positive). **B, C**, Slices with more than 50 ChrimsonR-CA3 neurons. Note the cFos positive CA1 neurons in addition to the CheRiff-CA1 neurons. Scale bars 100 µm.

Following two-color light stimulation inside the incubator, the organotypic cultures remained untouched for three full days. After this incubation period, they were placed in the recording chamber of the patch-clamp setup (Fig. 2A). To determine whether oSTDP had induced long-term changes in synaptic strength, EPSCs were recorded from CA1 neurons in response to stimulating ChrimsonR-CA3 neurons with 1 ms flashes of 594 nm light (Fig. 4A). In these and all following experiments, the experimenter was blind to the stimulation pattern, and stimulated (or otherwise treated) slice cultures were always interspersed with each other and non-paired controls. To obtain a relative measure of synaptic input strength, we compared the EPSC measured in each transfected CA1 neuron to the average of EPSCs recorded from several non-transfected CA1 neurons in the same field of view (Fig. 4B-C, see methods). There were no differences in input strength of ChrimsonR-CA3 to CheRiff-CA1 neurons in control non-paired slices (no light, postsynaptic only) or slices with mKate2-only CA1 neurons (Fig. 4D, p = 0.62 ANOVA). Thus, expression of the optogenetic tools did not affect synaptic connections, and we pooled the non-paired control experiments for statistical analysis (see below, Supplementary Table 1). Synapses between ChrimsonR-CA3 and CheRiff-CA1 neurons were significantly potentiated three days after causal pairing (Fig. 4E, +10 ms: p = 0.003, Dunnett’s). Unexpectedly, synapses were also potentiated (rather than depressed) 3 days after anti-causal pairing (Fig. 4E, -10 ms: p = 0.014, Dunnett’s). Still, the time interval between EPSPs and postsynaptic spikes was important, as input strength did not change when Δt was 50 ms (Fig. 4E, +50 ms: p = 0.49, Dunnett’s; -50 ms: p = 0.49, Dunnett’s). Thus, three days after oSTDP, a clear trace of the 5 Hz induction episode remained if CA3 and CA1 cells were activated synchronously (+/- 10 ms), but when activated asynchronously (+/- 50 ms).

**Figure 4.**
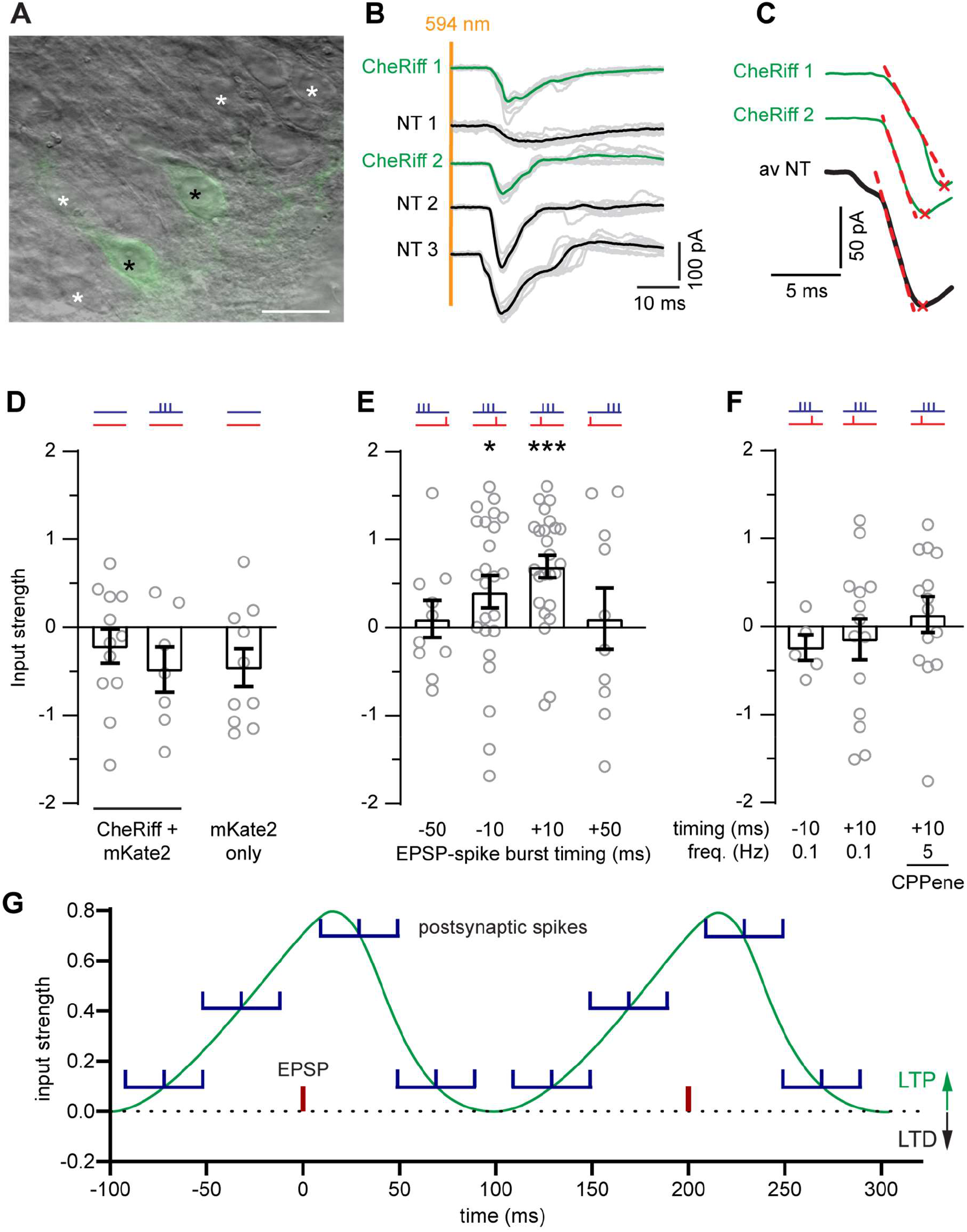
Input to Cheriff-CA1 neurons three days after optogenetic STDP. **A**, Dodt-contrast image (40x objective) of the CA1 region with overlaid epifluorescence image. Black asterisks: CheRiff-eGFP expressing CA1 pyramidal neurons; white asterisks: non-transfected CA1 pyramidal neurons suitable for recording. Scale bar 25 µm. **B**, Yellow light (1 ms, 594 nm) on ChrimsonR-CA3 neurons evoked excitatory postsynaptic currents (EPSCs) in CA1 neurons of a control (no oSTDP pairing) slice. EPSCs are recorded sequentially from CheRiff-CA1 pyramidal neurons (green, average of 10 grey individual EPSCs) and at least three non-transfected (NT) CA1 neurons (black average of 10 grey individual EPSCs). **C**, Automatically-detected EPSC peak (red x) and slope (dashed red line, 20-60% peak) from individual CheRiff-CA1 neurons and the average of NT neurons. **D-G**, Red and violet ticks indicate pre-(red) and postsynaptic (violet) light stimulation. **D**, Normalized input strength of CheRiff-CA1 neurons recorded from nonpaired control (left: no light stimulation; right: postsynaptic stimulation only) slices and mKate2-CA1 neurons three days later. n = 12; 7; 10 (left to right). **E**, Normalized input strength of CheRiff-CA1 neurons 3 days after 300 pairings of single presynaptic and 3 postsynaptic spikes at 5 Hz. During anti-causal pairing the last postsynaptic spike occurred -50 or -10 ms before the EPSP. During causal pairing the first postsynaptic spike occurred +10 or +50 ms after the EPSP. n = 10; 24; 25; 10 (left to right). *p < 0.05, ***p < 0.001. **F**, Same as E, but in groups 1 and 2, the pairing frequency was reduced to 0.1 Hz (360 pairings in 1 hour). Group 3: The NMDA receptor antagonist CPPene (1 µM) was in the culture medium during causal (+10 ms pairing 300x at 5 Hz. n = 5; 14; 14 (left to right). **G**, Mean input strength (data from E) as a function of timing between EPSPs (red) and postsynaptic spike bursts (violet, at mean) at 5 Hz repetition frequency. Two complete cycles are illustrated. No tLTD window was observed.

It has been reported that tLTP is associated with an immediate increase in postsynaptic excitability ^26– 28^. We tested for such changes three days after oSTDP, but found no differences in passive electrical parameters or excitability (number of action potentials in response to current injection) between the CheRiff-CA1 and corresponding non-transfected CA1 neurons (Supplementary Fig. 1). Thus, late tLTP reflects a change in synaptic strength rather than altered postsynaptic excitability.

### Late tLTP is frequency and NMDA receptor-dependent

We further investigated the requirements for induction of late tLTP. Previous studies have reported a strong dependence on pairing frequency ^21^. Likewise, reducing the pairing frequency from 5 Hz to 0.1 Hz prevented late tLTP in our experiments (0.1 Hz, -10 ms: p = 1; +10 ms: p = 0.87, Dunnett’s, Fig. 4F). Furthermore, as for classical STDP ^8,9,19,20,23^, late tLTP was abolished when NMDA receptors were blocked during pairing (+CPP: p = 0.23, Dunnett’s, Fig. 4F).

### The late appearance of tLTP after anti-causal pairing is activity-dependent

While optogenetic tLTD was observed 20 min after induction in patch-clamped neurons, we instead observed tLTP 3 days after in-incubator anti-causal oSTDP induction (Fig. 4G). We tested whether late tLTD might be uncovered by changing the number of anti-causal pairings ^21^. However, 100 to 500 pairings induced potentiation, while 30 repetitions were too few (30 rep.: p = 0.98; 100 rep.: p = 0.03; 300 rep.: p = 0.02; 500 rep.: p = 0.03, Dunnett’s, Fig. 5A). Could it be that STDP switches sign with time? We examined the outcome 2-3 hours after in-incubator anti-causal oSTDP. At this intermediate time point there was no significant change in input strength (Fig. 5B). We reasoned that after oSTDP, circuits containing paired synapses might develop higher levels of spontaneous spiking, increasing re-activation of the synapses between the paired CA3 and CA1 neurons thus leading to tLTP after anti-causal pairing ^28,29^. Accordingly, we globally suppressed activity starting 3-4 hours after anti-causal (−10 ms) pairing for two days and recorded from the treated slices 3 days later. Late tLTP after anti-causal pairing was abolished by this treatment (−10 ms, TTX: p = 1, Dunnett’s, Fig. 5B). These results suggest that the synaptic memory of a short episode of coincident activity is actively modified in the circuit.

**Figure 5:**
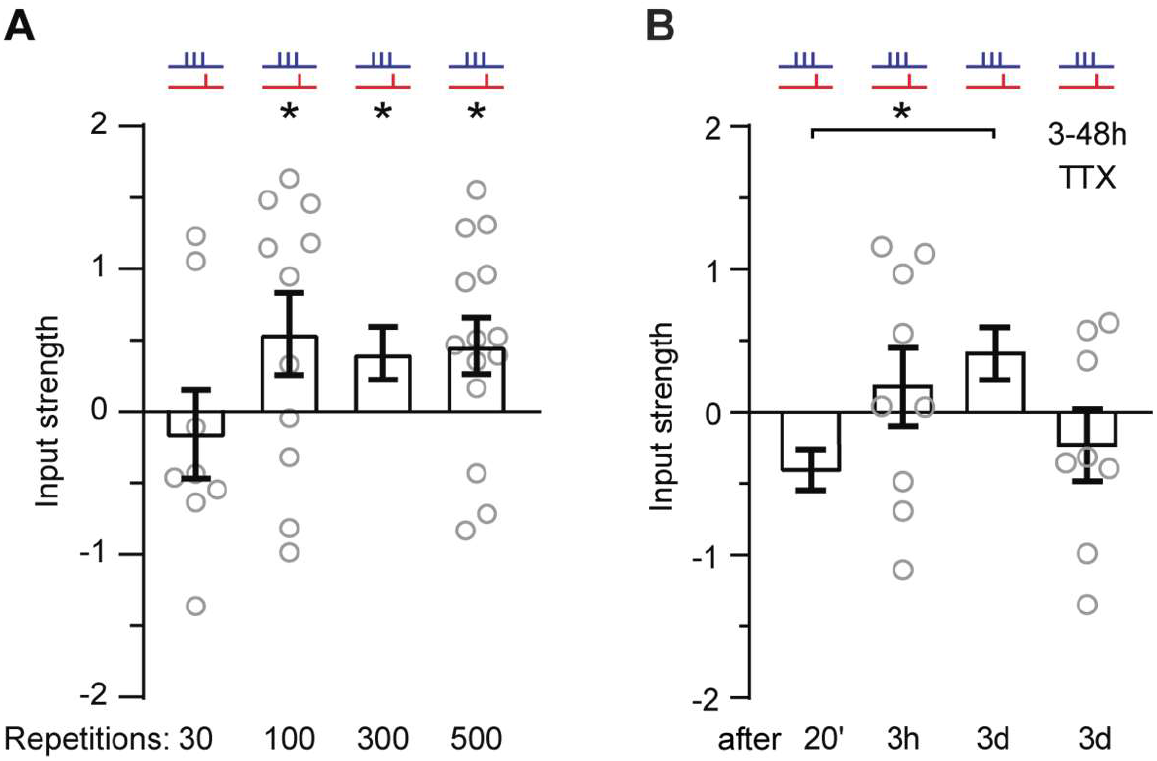
Late tLTP after anti-causal pairing is time and activity dependent. **A**, Input strength 3 days after anti-causal (−10 ms) pairing using different numbers of repetitions n = 8; 11; 24; 14 (left to right). The 300 repetition group is replotted from Fig.4E. * p < 0.05 mean ± SEM. **B**, Input strength at varying times after anti-causal pairing and when spiking was blocked with tetrodotoxin (TTX, 1 µM) from 3-4 hours until 1 day before assessment. n = 11, 9, 24, 8 (left to right). * p < 0.05 mean ± SEM The 20 minute and 3 day data are re-plotted from Fig. 2F and Fig. 4E.

## Discussion

Previously, the requirement of impaling the postsynaptic neuron with a sharp electrode or using whole-cell or perforated patch recordings to induce STDP has made it impossible to assess the late (i.e. three days later) effects of STDP. In the present study, we demonstrate: 1) that the opsins CheRiff and ChrimsonR can be used to independently spike CA3 and CA1 neurons, 2) that theta frequency optogenetic pairing in causal sequence (first CA3, then CA1) induces tLTP, whereas anti-causal pairing induces tLTD during patch-clamp recordings, and 3) that the long-term consequence of oSTDP is a symmetric potentiation-only window at short timing intervals (± 10 ms) and no potentiation at longer intervals (± 50 ms). Thus, we have directly demonstrated something long assumed to be true, that information about the coincidence of paired spiking is stored for an extended period in the strength of the connections between those specific neurons. As we normalized the strength of inputs onto the paired postsynaptic neurons to the input onto neighboring (unpaired) neurons, global changes during the three-day incubation period, would not affect our results. Similarly, increasing expression of presynaptic ChrimsonR would equally affect EPSC slopes in paired and neighboring NT neurons. Underlying late tLTP may be strengthening of existing synapses, an increase in the number of synapses between the paired neurons, or a combination of both. We cannot at present distinguish between these possibilities: Our method includes two different fluorescent labels in pre- and postsynaptic neurons, but to unequivocally identify and count synapses with light microscopy, additional functional information is required, e.g. activity-dependent labeling ^30^.

Since during our short-term patch-clamp recordings, oSTDP reproduced canonical asymmetric STDP with tLTD and tLTP after anti- and causal pairing respectively ^8–10,31,32^, we surmise that optically- and electrically-induced spiking are not fundamentally different. Also, late tLTP depends on NMDA receptors as is well established for electrode-induced STDP ^19^, strongly suggesting that elevated intracellular [Ca^2+^] in the postsynaptic neuron is essential. A further similarity of late oSTDP to electrode-induced STDP is the strong frequency-dependence ^21^: At a pairing repetition rate of 5 Hz, late tLTP is observed, but not when the pairing rate is reduced to 0.1 Hz. As in previous STDP studies, there is no potentiation at spike intervals of ±50 ms. When postsynaptic spikes occur about 50 ms after EPSPs, GABA_A_ currents from feed-forward inhibition are strongly active in CA1 neurons ^33^ and may reduce postsynaptic calcium influx, as has been demonstrated for back-propagating action potentials ^34^. Even with inhibition blocked, single spine synaptic calcium transients are reduced if paired bAPs are delayed by 40 ms compared with shorter delays ^22^. Postsynaptic AMPA currents have decayed and no longer contribute to the bAP-induced relief of NMDAR Mg^2+^ block. Likewise, postsynaptic spikes that precede EPSPs by more than a few ms will not contribute to the single-spine Ca^2+^ transients.

As optical anti-causal pairing induced tLTD during patch-clamp recordings, we were very surprised by the lack of late tLTD three days after oSTDP induction. While we cannot exclude that somewhere in the vast parameter space a protocol for late tLTD may exist, none of the modified protocols we tested (increasing or decreasing the number of anti-causal pairing repetitions, changing the pairing interval, decreasing the pairing frequency) reduced synaptic input three days after induction. Nor is tLTD apparent at the earliest time window (3 hours) feasible after noninvasive oSTDP induction. Interestingly, Pang et al. ^35^, who used extracellular electrodes to induce and record STDP failed to observe LTD at early (< 30 min) time points. During their 4-hour long recordings, LTP persisted for timings from -10 ms to +20 ms and LTD slowly appeared only for the anti-causal pairing interval of -20 ms. Both their study and the present work suggest that STDP has hitherto unappreciated slow dynamics that can include a change in sign.

When we disrupted spontaneous activity in the slice cultures 3 hours after oSTDP, the ‘synaptic memory’ of oSTDP was erased. This led us to hypothesize that the endogenous activity in the circuit becomes biased towards ‘replaying’ the optogenetically-induced sequences, reactivating the paired synapses. Interestingly, maintenance and consolidation of memories in vivo also requires ongoing activity and NMDA receptors ^36,37^. Transient knock-out of NMDA receptors wipes out previously encoded memory, but does not prevent the future acquisition of new memories ^36^. Indeed, the notion that memory consolidation requires replay, for instance during sleep, is well supported ^38,39^. Directly testing this hypothesis will require continuous monitoring of circuit activity with synaptic resolution, ideally simultaneously at many synapses over several days.

That the early effects of oSTDP replicated the classical asymmetric STDP window strongly argues against optogenetic artifacts being responsible for the difference in early and late effects. As channelrhodopsins are distributed in the plasma membrane, it could be argued that optical generation of spikes is actually more physiological than somatic current injection: Natural spikes arise from summation of active excitatory synapses distributed over the dendritic arbor, not from spontaneous depolarization of the soma. Dendritic depolarization rather than actual spiking of the postsynaptic neurons is likely important for synaptic plasticity: Experiments pairing mossy fiber stimulation with CA3-CA3 synaptic inputs induces tLTP or tLTD of the CA3-CA3 synapses without spiking of the postsynaptic CA3 neuron^40^. Likewise, at CA3-CA1 Schaffer collateral synapses, tLTP can occur without postsynaptic spikes ^41^. Therefore, if CheRiff-CA1 cells occasionally missed spiking and the dendrites only depolarized in response to the violet light flashes, synaptic plasticity would still be expected.

While the majority of STDP studies report both potentiation and depression, ‘LTP-only’ STDP windows have been observed at human hippocampal, human and rat neocortical synapses ^42,43^ and in the mouse hippocampus ^21^, most commonly in the presence of increased dopamine ^32,43^. If presynaptic activity is paired with prolonged postsynaptic bursts (plateau potentials), the timing window for Schaffer collateral potentiation can be extended to several seconds in the causal and anti-causal direction ^44^. While there are many possible explanations for these discrepant outcomes of early oSTDP, we speculate that if the neurons are left intact and postsynaptic strength is assessed after several days, the outcome of repeated coincident activity at Schaffer collateral synapses may always be potentiation at short pairing intervals. What we do not believe is that there is no LTD. Heterosynaptic LTD or generalized downscaling processes are important mechanisms to conserve total synaptic weight and prevent runaway potentiation^45^. Although unlikely, we cannot exclude that the non-paired synapses onto the NT-CA1 neurons and the ± 50 ms paired synapses underwent LTD relative to the ± 10 ms paired synapses as we assessed relative synaptic strength. To put the relationship between synaptic plasticity and memory storage onto a firm empirical basis, we need to investigate how neuronal activity modifies synaptic connections though STDP-like processes over extended periods of time.

## Acknowledgments

We would like to thank Mark van Rossum for critically reading the manuscript. Iris Ohmert, Sabine Graf, Torsten Renz, Dirk Lubrich, Lennart Sobirey, Paul Lamothe-Molina and Ingke Braren (UKE Vector facility) provided important technical support. CheRiff was a gift from Adam Cohen, ChrimsonR was a gift from Edward Boyden. The first spectral characterization of ChrimsonR-expressing neurons in slice cultures was performed by Katie Ferguson and Longzhi Tan during the 2014 Neurobiology Course at MBL in Woods Hole. Funding was received from: the Landesforschungsförderung Hamburg; the Deutsche Forschungsgemeinschaft (DFG) grants SPP 1665 220176618, SPP1926 315380903, SFB 936 178316478, SFB 1328 335447717, FOR 2419 278170285, Emmy Noether 259979908 and the European Research Council (ERC-StG 714762).

## Author Contributions

**Conceptualization**, CEG, TGO, JSW; **Investigation**, MA, BvB, CEG; **Writing – Original Draft**, CEG, TGO, MA; **Writing – Review and Editing**, CEG, TGO, MA, BvB, JSW, MM; **Technical expertise**, TGO, CEG, MM; **Visualization**, MA, CEG, TGO; **Supervision**, CEG, TGO; **Project Funding Acquisition**, CEG, JSW. All authors approved the author list and final version.

## Materials and Methods

### Rat organotypic hippocampal slice cultures preparation and transfection

Wistar rats were housed and bred at the University Medical Center Hamburg-Eppendorf (UKE) animal facility and sacrificed according to German Law (Tierschutzgesetz der Bundesrepublik Deutschland, TierSchG) with approval from the Behörde für Justiz und Verbraucherschutz (BJV)-Lebensmittelsicherheit und Veterinärwesen, Hamburg and the animal care committee of the UKE. The procedure for preparing organotypic cultures was modified from Stoppini et al. ^46^, using media without antibiotics ^47^. Cultures were transduced at 10-11 days in vitro (DIV) with local injection^48^ into CA3 of AAV2.Rh10-syn-ChrimsonR-tdTomato 7.22×10^13^ vg ml^-1^ (packaged at the UKE Vector Facility, plasmid a gift from Edward Boyden). On the same day CA1 neurons were transfected with pAAV-hsyn-CheRiff-eGFP (0.5 ng µl^-1^) and pCI-syn-mKate2 (10 ng µl^-1^) using single cell electroporation ^49^. After opsins were expressing, great care was taken to protect the cultures from wavelengths of light below 560 nm to prevent unintentional co-activation of ChrimsonR and CheRiff-expressing neurons. Handling and normal medium changes (2/3, twice per week) were performed under dim yellow light (Osram LUMILUX CHIP control T8).

### CheRiff and ChrimsonR characterization

For characterizing the responses to multiple wavelengths, we used a pE-4000 CoolLED (CoolLED Ltd., s/n CP0180) with an additional filter for 550 nm (555 ± 20 nm). Light intensities in the specimen plane were measured using a 918D-ST-UV sensor and a 1936R power meter (Newport). Light flashes (1 ms, low to high intensity) were delivered every 20 s with one minute between wavelength sequences. Whole cell currents were recorded (see electrophysiology below) with synaptic transmission and action potentials blocked by CPPene (10 µM, Tocris bioscience; 1265), NBQX (10 μM, Tocris bioscience; 1044), picrotoxin (100 µM, Sigma; P1675-1G) and tetrodotoxin (1 µM, HelloBio; HB1035). Seven CA1 (CheRiff) and eleven CA3 (ChrimsonR) neurons were recorded from to obtain n = 4 responses at each wavelength/intensity combination. Recordings were performed in the recording medium described below (see optogenetic induction of spike-timing dependent plasticity (oSTDP) during whole-cell recordings).

### Combined electrophysiology and optogenetic stimulation

The setup is based on an Olympus BX61WI microscope fitted with a Dodt contrast, epifluorescence and CCD-camera (DMK23U274, The Imaging Source) on a dual camera port (see Fig. 2A). For light stimulation through the objective, a LED combiner (Mightex Systems, wavelengths 625, 530, 470, 405 nm) was coupled via a 1 mm multimode fiber and a collimator (Thorlabs) mounted on one of the camera ports. A 40x water immersion objective (Plan-Apochromat 1.0 NA DIC VIS-IR, Zeiss, illuminated field ø 557 µm) or a 10x water immersion objective (UMPlanFL 0.30 NA W, Olympus, illuminated field ø 2.65 mm) were used. The light intensities through the objectives were measured in the specimen plane using a calibrated power meter (LaserCheck, Coherent or 918D-ST-UV sensor / 1936R power meter, Newport). To locally activate the CA3 neurons, the orange (594) laser beam of an Omicron Light Hub was coupled into an optical fiber fitted with a collimator (Thorlabs) mounted on a lockable swing arm. This construction allowed us to point the laser beam at different angles through the center of the back aperture of the oil immersion condenser (NA 1.4, Olympus) to locally illuminate neurons in CA3 away (∼1.5 mm) from the optical axis, with high intensity and little light scattering (Fig. 2A, B). After final laser positioning, the stage was not moved and CA1 neurons within the single field of view were recorded from.

Whole-cell patch-clamp recordings were performed using an Axopatch 200B (Axon Instruments, Inc.) amplifier, National Instruments A/D boards and Ephus software ^50^. EPSCs (Fig. 4) and cellular parameters (Supplemental Fig. 1) were measured at 30°C (in-line heater, Warner Instruments) in artificial cerebral spinal fluid containing: NaCl (119 mM, Sigma; S5886-500G), NaHCO_3_ (26.2 mM, Sigma; S5761-500G), D-glucose (11 mM, Sigma; G7528-250G), KCl (2.5 mM, Fluka; 60121-1L), NaH_2_PO_4_ (1 mM, Sigma; S5011-100G), MgCl_2_ (4 mM, Fluka; 63020-1L), CaCl_2_ (4 mM, Honeywell; 21114-1L), pH 7.4, ∼310 mOsm kg^-1^, saturated with humidified 95% O_2_ / 5% CO_2_. The patch electrodes (3-4 MΩ, World Precision Instruments; 1B150F-3) were filled with intracellular solution containing: K-gluconate (135 mM, Sigma; G4500-100G), EGTA (0.2 mM, Sigma-Aldrich; E0396-10G), HEPES (10 mM, Sigma; H4034-100G), MgCl_2_ (4 mM, Fluka; 63020-1L), Na_2_-ATP (4 mM, Aldrich; A26209-1G), Na-GTP (0.4 mM, Sigma; G8877-100MG), Na_2_-phosphocreatine (10 mM, Sigma; P7936-1G), ascorbate (3 mM, Sigma; A5960-100G), pH 7.2, 295 mOsm kg^-1^. The liquid junction potential was measured (−14.1 to -14.4 mV) and corrected. CA1 pyramidal neurons were patched under the 40x water immersion objective and voltage-clamped at -70 mV (LJP corrected). A test pulse (−5 mV, 200 ms) was applied at the start of every sweep. The series resistance (Rs) was less than 20 MΩ, did not change more than 30% and was not compensated in voltage clamp. In current clamp, we used bridge balance compensation.

### Optogenetic induction of spike-timing dependent plasticity (oSTDP) during whole-cell recordings

Eight to eleven days after virus injection and electroporation of ChrimsonR and CheRiff (18-22 DIV), the slices were transferred to the electrophysiology setup and perfused with slice culture medium containing (for 500 ml): 494 ml Minimal Essential Medium (Sigma M7278), 1 mM L-glutamine (Gibco 25030-024), 0.01 mg ml^−1^ insulin (Sigma I6634), 1.45 ml 5 M NaCl (S5150 Sigma), 2 mM MgSO_4_ (Fluka 63126), 1.44 mM CaCl_2_ (Fluka 21114), 0.00125% ascorbic acid (Fluka 11140), 13 mM D-glucose (Fluka 49152), supplemented with D-serine (30 μM, Tocris Bioscience; 0226) and diluted slightly to give ∼308 mOsm k^-1^ (33°C, saturated with humidified 95% O_2_ / 5% CO_2_). The LJP was corrected (−15.3 mV). The medium (50 ml) was recycled during the experiment and changed every 4 hours or every 4 slices, whichever came first. With the 594 nm laser positioned to stimulate ChrimsonR-CA3 neurons, a CA1 neuron was patched and EPSCs were recorded (every 20s) at -65.3 mV (LJP corrected) where inhibitory currents were clearly outward. The laser intensity was set to evoke EPSCs of approximately -100 pA (range -50 pA to -200 pA). Baseline EPSCs were recorded for up to 5 minutes after break-in. The 10x objective was carefully moved into position to maximize the area of the slice illuminated and the amplifier was switched to current clamp to allow spiking of the CA1 neuron during paired optogenetic stimulation. To induce plasticity, stimulation of ChrimsonR-CA3 neurons (300, 2 ms pulses at 5 Hz (594 nm laser at baseline intensity) plus simultaneous red LED stimulation through the objective (625 nm, 2 mW mm^-2^)) was paired with bursts of 3 violet LED stimuli in CA1 (405 nm, 2 ms, 50 Hz, 1 mW mm^-2^) to activate CheRiff-CA1 neurons. After pairing CA3 and CA1 stimulation, EPSCs (0.05 Hz orange laser flashes) were recorded for additional 25 minutes. Pairing was always completed within 10 minutes of break-in.

### In-incubator oSTDP, read-out and analysis

We waited 7-10 days (17-21 DIV) to ensure stable expression of ChrimsonR and CheRiff. The closed petri dish containing the centered slice culture was placed in an LED illumination tower constructed from 30 mm optical bench parts (Thorlabs) situated inside a dedicated incubator. The tower contained an injection-molded reflector (Roithner LaserTechnik, 10034) to collimate the 625 nm LED (Cree XP-E red) and an aspheric condenser lens (Thorlabs ACL2520U-A) to collimate the 405 nm LED (Roschwege Star-UV405-03-00-00). The LEDs were powered and controlled from outside the incubator by a two-channel Grass S8800 stimulator, two constant-current drivers (RECOM RCD-24-1.20) and a timer. The LED power was adjusted to give 1 mW mm^-2^ for 405 nm and 8 mW mm^-2^ for 625 nm inside the petri dish in the specimen plane. ‘Non-paired’ cultures were handled identically, but no light stimulation was given. After pairing, each petri dish was marked with a letter code and date/time of pairing. Where indicated, CPPene (1 µM, Fig. 4H) was added to the slice culture medium the day before or (Fig. 4I) 3-4 hours after pairing the medium was changed to medium containing TTX (1 µM) to stop all spiking. Two days later, the slices were washed with fresh medium and returned to the incubator until the following day when they were recorded from.

Three days or one hour after in-incubator stimulation, the slices were transferred by a different blinded investigator to the electrophysiology setup with on/off axis stimulation and perfused with ACSF at 30 °C as described above. Whole-cell patch-clamp recordings were sequentially made from 3 to 4 NT-CA1 pyramidal neurons and 1 to 5 CheRiff-CA1 pyramidal neurons in a pseudo-random order without moving the stage to ensure constant CA3 stimulation (holding potential -70 mV to ensure separation between EPSCs and IPSCs). While still in cell-attached mode, CA1 neurons were illuminated with a single 405 nm light flash (1 ms, 1 mW mm^-2^). None of the 255 NT-CA1 neurons spiked. Of 224 fluorescent CheRiff-CA1 neurons, 96% spiked in response to the single violet flash, 202 fired a single spike (90.2%) and 13 fired 2-3 spikes (5.8%). The recordings were immediately discontinued from the remaining 9 neurons that did not spike (4%).

After break-in, ChrimsonR-CA3 neurons were stimulated through the condenser (594 nm, 20 s interval) to evoke EPSCs in the CA1 neurons. The EPSC onset was typically 6 to 8 ms after the stimulation. If the delay to the EPSC onset was 13 ms or longer, connections were assumed to be polysynaptic and not analyzed. Ten EPSCs were recorded at 3 laser intensities from each CA1 neuron (when spontaneous events obscured the EPSCs, extra sweeps were collected). The middle intensity was set to evoke an EPSC of about -100 to -300 pA in the first recording from a given slice. The same 3 intensities were used for all neurons in a given slice, but only one intensity was used for analysis. The selected intensity evoked the most similar EPSC slopes that were at least 20 pA in three non-transfected cells (usually the lowest laser intensity).

At the end of each recording, we injected a series of current steps to assess cellular excitability. Cells without pyramidal-like firing patterns (i.e. high AP frequency, large amplitude after-hyperpolarization) were eliminated from the analysis. Individual sweeps with spontaneous EPSCs/IPSCs preceding the stimulation by less than 10 ms were not analyzed. If fewer than 4 sweeps remained at the selected laser intensity, the recording was discarded. A custom MATLAB routine averaged the EPSCs, detected the peak and the 20-60% slope (Fig. 4 C). If the time to peak varied by more than 5 ms between neurons, we suspected a mixture of mono- and polysynaptic responses and the slice was excluded.

### Immunohistochemistry

An hour after stimulation, the slices were fixed (30 minutes, ice-cold 4% paraformaldehyde in PBS). The slices were washed in PBS (3 x 10 min), blocked (2 hours 5% donor horse serum, 0.3 % TritonTM X-100 in PBS at room temperature) and then incubated overnight with the primary antibody (4° C, rabbit anti-cFos, Santa Cruz Biotechnology, Inc., sc-52, 1:500) in carrier solution (1 % Bovine serum albumin, 0.3 % TritonTM X-100 in PBS). On the next day, the slices were washed 3 times in PBS and incubated at room temperature for 2 hours with a secondary antibody (Alexa Fluor 568 goat anti-rabbit, Life Technologies, A11011) at 1:1000 in carrier solution (same as above) and afterwards washed again (3 x 10 min) in PBS and mounted with a Shandon Immu-Mount (Thermo Scientific; 9990402).

### Confocal microscopy

An Olympus F1000 confocal microscope with 20x objective (UPLSAPO 0.85 NA, Olympus) and filter sets for GFP, RFP (tdTomato) and Alexa 568 was used for immunocytochemistry. Z-series images were obtained using 3 µm z-steps at a 1024×1024-pixel resolution scanning at 12.5 µs per pixel. The imaging parameters for the red and green channel were kept constant throughout all experiments. Fiji/ImageJ ^51^ was used to generate Z-projections and to overlay channels.

Quantification and statistical analysis

### Experimental design

Except for data presented in Figures 1 and 2, all experiments and analysis were performed blind to the in-incubator optical stimulation or any drug treatments. Unblinding only occurred after initial analyses and any decision to include/exclude recordings had been taken. To ensure blinding, we always interspersed different treatment conditions with control (non-paired) slices (Fig. 4D) and ±10 ms groups (Fig. 4E).

### Normalization

Input strength was defined according to the formula:

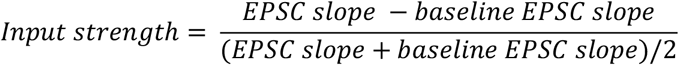

where (Fig. 2) EPSC slope is the average from 20-25 minutes after oSTDP pairing and baseline the average EPSC slope (∼5 minutes) before pairing, or (Fig. 4 & 5) ‘baseline’ is the average EPSC slope of all non-transfected (NT-CA1) neurons from the same slice.

### Statistical analysis

GraphPad Prism 9 was used for statistical analysis and to generate plots. All data is represented as mean ± standard error of the mean (SEM). Statistical significance was assumed for p < 0.05. For details of the statistical analysis see Supplementary Table 1.

## Supplemental Information

**Supplementary Figure 1.**
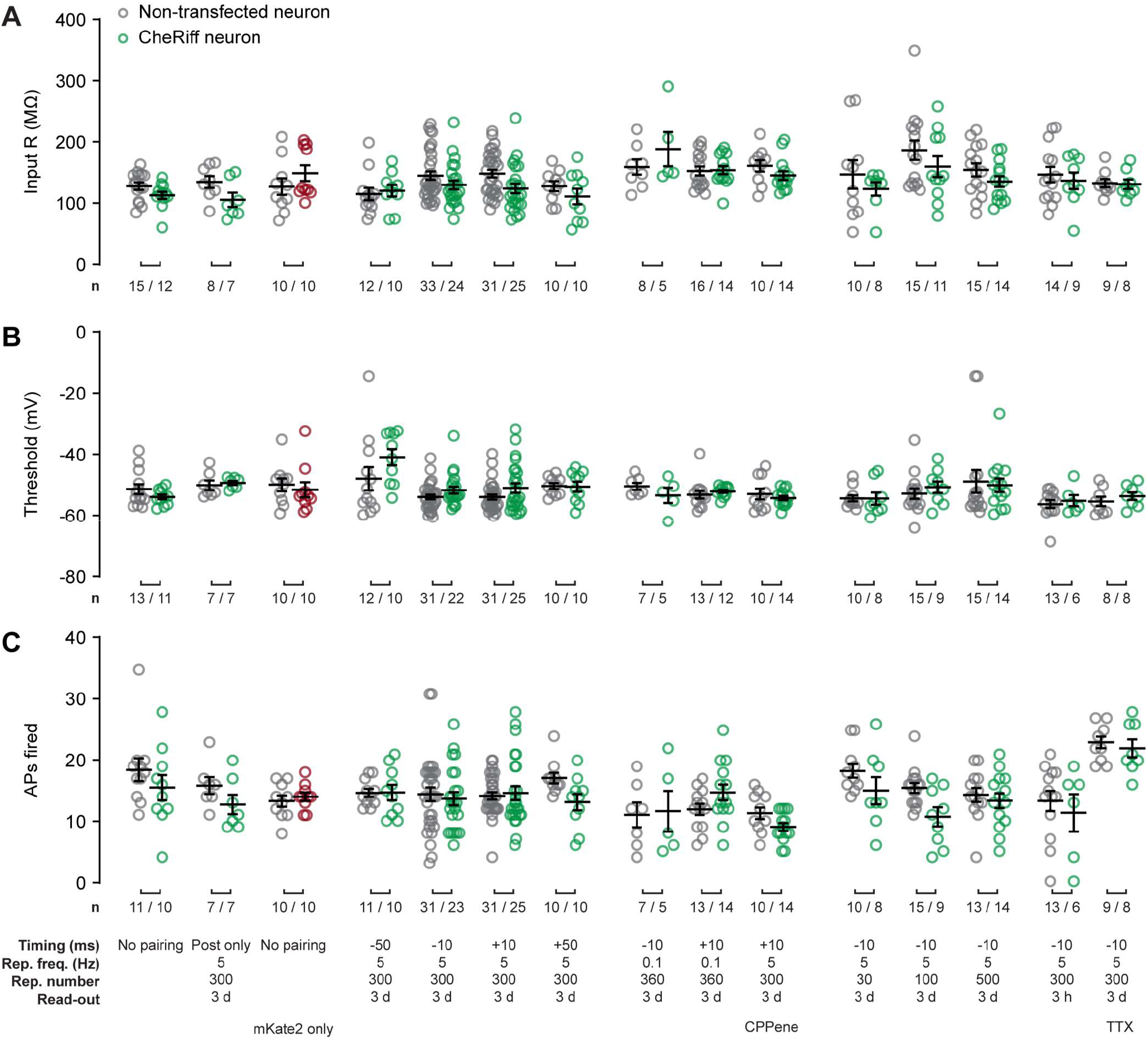
Comparison of passive and active cell parameters of CheRiff-CA1 and neighboring NT-CA1 neurons from oSTDP experiments in Figure 4. **A**, Input resistance of all NT-CA1 and CheRiff-CA1 neurons. **B**, Action potential threshold. **C**, Numbers of action potentials fired in response to a 400 pA current step. There were no significant differences between NT and CheRiff-CA1 neurons in any of the treatment groups (Kolmogorov-Smirnov). CPPene: 1 µM CPPene during oSTDP; TTX: 1 µM tetrodotoxin 4-48 h after oSTDP. n number of NT-CA1 / CheRiff-CA1 neurons. Plotted are individual data points, mean ± SEM. As the current clamp measurements (**B** and **C**) were performed last, they were missed in some recordings.

**Supplementary Table 1:**
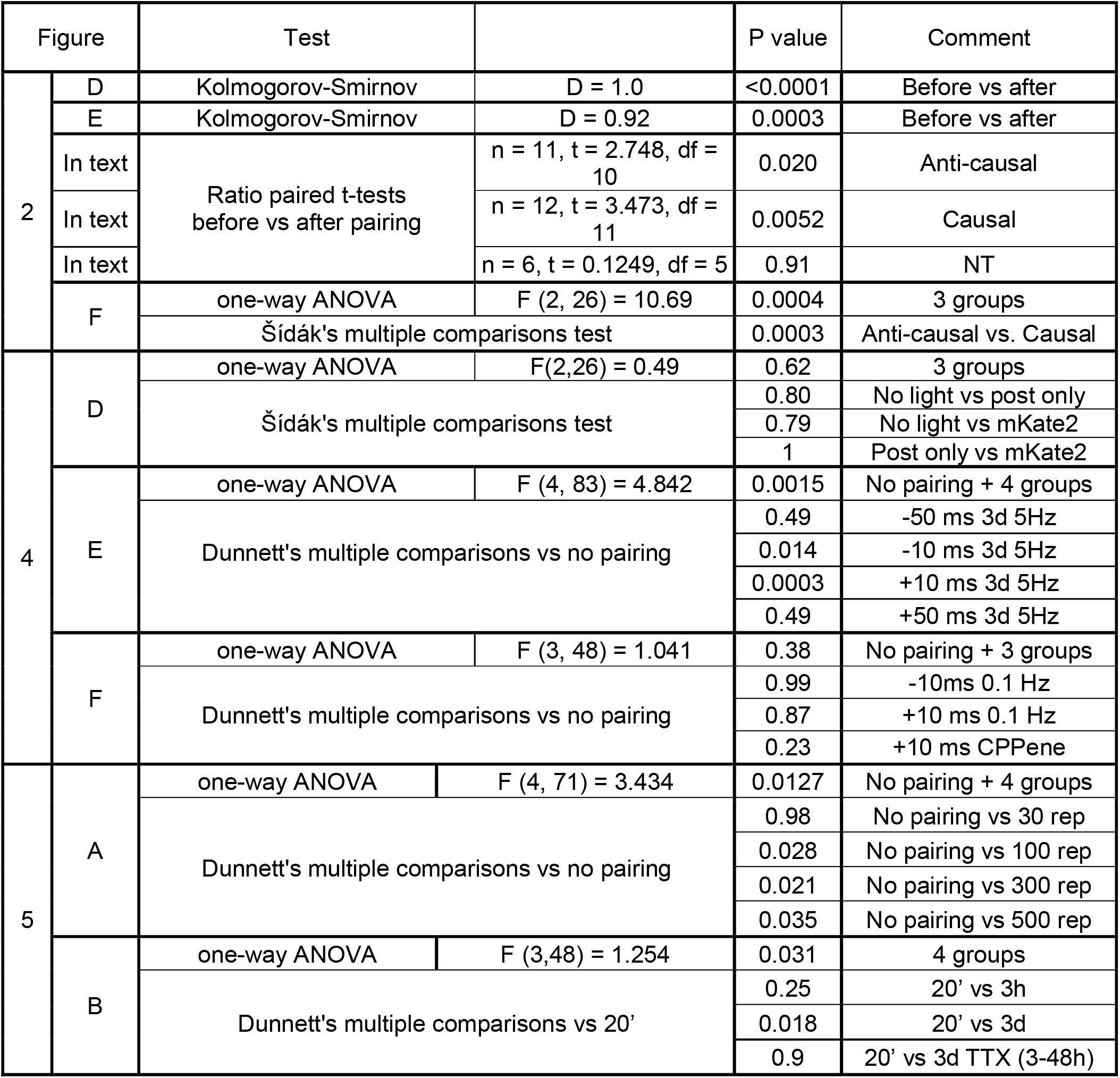
Summary of statistical analyses. All analyses were performed using GraphPad Prism

## References

1. Song, S., Miller, K.D., and Abbott, L.F. (2000). Competitive Hebbian learning through spike-timing-dependent synaptic plasticity. Nat. Neurosci. 3, 919–926.

2. Gerstner, W., Kempter, R., van Hemmen, J.L., and Wagner, H. (1996). A neuronal learning rule for sub-millisecond temporal coding. Nature 383, 76–81.

3. Costa, R.P., Froemke, R.C., Sjöström, P.J., and van Rossum, M.C.W. (2015). Unified pre-and postsynaptic long-term plasticity enables reliable and flexible learning. Elife 4, e09457.

4. Masquelier, T., Guyonneau, R., and Thorpe, S.J. (2009). Competitive STDP-Based Spike Pattern Learning. Neural Comput. 21, 1259–1276.

5. Serrano-Gotarredona, T., Masquelier, T., Prodromakis, T., Indiveri, G., and Linares-Barranco, B. (2013). STDP and STDP variations with memristors for spiking neuromorphic learning systems. Front. Neurosci. 7, 2.

6. Kheradpisheh, S.R., Ganjtabesh, M., Thorpe, S.J., and Masquelier, T. (2018). STDP-based spiking deep convolutional neural networks for object recognition. Neural Networks 99, 56–67.

7. Bing, Z., Meschede, C., Chen, G., Knoll, A., and Huang, K. (2020). Indirect and direct training of spiking neural networks for end-to-end control of a lane-keeping vehicle. Neural Netw. 121, 21– 36.

8. Debanne, D., Gähwiler, B.H., and Thompson, S.M. (1998). Long-term synaptic plasticity between pairs of individual CA3 pyramidal cells in rat hippocampal slice cultures. J. Physiol. 507, 237–247.

9. Bi, G., and Poo, M. (1998). Synaptic Modifications in Cultured Hippocampal Neurons: Dependence on Spike Timing, Synaptic Strength, and Postsynaptic Cell Type. J. Neurosci. 18, 10464–10472.

10. Markram, H. (1997). Regulation of Synaptic Efficacy by Coincidence of Postsynaptic APs and EPSPs. Science (80-.). 275, 213–215.

11. Brzosko, Z., Mierau, S.B., and Paulsen, O. (2019). Neuromodulation of Spike-Timing-Dependent Plasticity: Past, Present, and Future. Neuron 103, 563–581.

12. Foncelle, A., Mendes, A., Jędrzejewska-Szmek, J., Valtcheva, S., Berry, H., Blackwell, K.T., and Venance, L. (2018). Modulation of Spike-Timing Dependent Plasticity: Towards the Inclusion of a Third Factor in Computational Models. Front. Comput. Neurosci. 12, 49.

13. Nagel, G., Szellas, T., Huhn, W., Kateriya, S., Adeishvili, N., Berthold, P., Ollig, D., Hegemann, P., and Bamberg, E. (2003). Channelrhodopsin-2, a directly light-gated cation-selective membrane channel. Proc. Natl. Acad. Sci. 100, 13940–13945.

14. Boyden, E.S., Zhang, F., Bamberg, E., Nagel, G., and Deisseroth, K. (2005). Millisecond-timescale, genetically targeted optical control of neural activity. Nat. Neurosci. 8, 1263–1268.

15. Huber, D., Petreanu, L., Ghitani, N., Ranade, S., Hromádka, T., Mainen, Z., and Svoboda, K. (2008). Sparse optical microstimulation in barrel cortex drives learned behaviour in freely moving mice. Nature 451, 61–4.

16. Adamantidis, A.R., Zhang, F., Aravanis, A.M., Deisseroth, K., and de Lecea, L. (2007). Neural substrates of awakening probed with optogenetic control of hypocretin neurons. Nature 450, 420– 424.

17. Klapoetke, N.C., Murata, Y., Kim, S.S., Pulver, S.R., Birdsey-Benson, A., Cho, Y.K., Morimoto, T.K., Chuong, A.S., Carpenter, E.J., Tian, Z., et al. (2014). Independent optical excitation of distinct neural populations. Nat. Methods 11, 338–46.

18. Hochbaum, D.R., Zhao, Y., Farhi, S.L., Klapoetke, N., Werley, C.A., Kapoor, V., Zou, P., Kralj, J.M., Maclaurin, D., Smedemark-Margulies, N., et al. (2014). All-optical electrophysiology in mammalian neurons using engineered microbial rhodopsins. Nat. Methods 11, 825–833.

19. Nevian, T., and Sakmann, B. (2006). Spine Ca2+ Signaling in Spike-Timing-Dependent Plasticity. J. Neurosci. 26, 11001–11013.

20. Tigaret, C.M., Olivo, V., Sadowski, J.H.L.P., Ashby, M.C., and Mellor, J.R. (2016). Coordinated activation of distinct Ca2+ sources and metabotropic glutamate receptors encodes Hebbian synaptic plasticity. Nat. Commun. 7, 10289.

21. Wittenberg, G.M., and Wang, S.S. (2006). Malleability of spike-timing-dependent plasticity at the CA3-CA1 synapse. J. Neurosci. 26, 6610–6617.

22. Holbro, N., Grunditz, A., Wiegert, J.S., and Oertner, T.G. (2010). AMPA receptors gate spine Ca(2+) transients and spike-timing-dependent potentiation. Proc. Natl. Acad. Sci. 107, 15975– 15980.

23. Edelmann, E., Cepeda-Prado, E., Franck, M., Lichtenecker, P., Brigadski, T., and Leßmann, V. (2015). Theta Burst Firing Recruits BDNF Release and Signaling in Postsynaptic CA1 Neurons in Spike-Timing-Dependent LTP. Neuron 86, 1041–1054.

24. Schoenenberger, P., Gerosa, D., and Oertner, T.G. (2009). Temporal control of immediate early gene induction by light. PLoS One 4, e8185.

25. Jaworski, J., Kalita, K., and Knapska, E. (2018). c-Fos and neuronal plasticity: the aftermath of Kaczmarek’s theory. Acta Neurobiol. Exp. (Wars). 78, 287–296.

26. Frick, A., Magee, J., and Johnston, D. (2004). LTP is accompanied by an enhanced local excitability of pyramidal neuron dendrites. Nat. Neurosci. 7, 126–135.

27. Debanne, D., and Poo, M.-M. (2010). Spike-timing dependent plasticity beyond synapse - pre-and post-synaptic plasticity of intrinsic neuronal excitability. Front. Synaptic Neurosci. 2, 21.

28. Debanne, D., Inglebert, Y., and Russier, M. (2019). Plasticity of intrinsic neuronal excitability. Curr. Opin. Neurobiol. 54, 73–82.

29. Sadowski, J.H.L.P., Jones, M.W., and Mellor, J.R. (2016). Sharp-Wave Ripples Orchestrate the Induction of Synaptic Plasticity during Reactivation of Place Cell Firing Patterns in the Hippocampus. Cell Rep. 14, 1916–1929.

30. Moeyaert, B., Holt, G., Madangopal, R., Perez-Alvarez, A., Fearey, B.C., Trojanowski, N.F., Ledderose, J., Zolnik, T.A., Das, A., Patel, D., et al. (2018). Improved methods for marking active neuron populations. Nat. Commun. 9, 4440.

31. Meredith, R.M., Floyer-Lea, A.M., and Paulsen, O. (2003). Maturation of long-term potentiation induction rules in rodent hippocampus: role of GABAergic inhibition. J. Neurosci. 23, 11142– 11146.

32. Zhang, J.-C., Lau, P.-M., and Bi, G.-Q. (2009). Gain in sensitivity and loss in temporal contrast of STDP by dopaminergic modulation at hippocampal synapses. Proc. Natl. Acad. Sci. U. S. A. 106, 13028–13033.

33. Samulack, D.D., and Lacaille, J.-C. (1993). Hyperpolarizing synaptic potentials evoked in CA1 pyramidal cells by glutamate stimulation of interneurons from the oriens/alveus border of rat hippocampal slices. II. sensitivity to GABA antagonists. Hippocampus 3, 345–358.

34. Marlin, J.J., and Carter, A.G. (2014). GABA-A receptor inhibition of local calcium signaling in spines and dendrites. J. Neurosci. 34, 15898–15911.

35. Pang, K.K.L., Sharma, M., Krishna-K., K., Behnisch, T., and Sajikumar, S. (2019). Long-term population spike-timing-dependent plasticity promotes synaptic tagging but not cross-tagging in rat hippocampal area CA1. Proc. Natl. Acad. Sci. 116, 5737–5746.

36. Cui, Z., Wang, H., Tan, Y., Zaia, K.A., Zhang, S., and Tsien, J.Z. (2004). Inducible and Reversible NR1 Knockout Reveals Crucial Role of the NMDA Receptor in Preserving Remote Memories in the Brain. Neuron 41, 781–793.

37. Shimizu, E., Tang, Y.P., Rampon, C., and Tsien, J.Z. (2000). NMDA receptor-dependent synaptic reinforcement as a crucial process for memory consolidation. Science (80-.). 290, 1170–1174.

38. Wilson, M.A., and McNaughton, B.L. (1994). Reactivation of hippocampal ensemble memories during sleep. Science 265, 676–9.

39. Squire, L.R., Genzel, L., Wixted, J.T., and Morris, R.G. (2015). Memory consolidation. Cold Spring Harb. Perspect. Biol. 7, a021766.

40. Brandalise, F., and Gerber, U. (2014). Mossy fiber-evoked subthreshold responses induce timing-dependent plasticity at hippocampal CA3 recurrent synapses. Proc. Natl. Acad. Sci. 111, 4303– 4308.

41. Hardie, J., and Spruston, N. (2009). Synaptic Depolarization Is More Effective Than Back-Propagating Action Potentials During Induction of Associative Long-Term Potentiation in Hippocampal Pyramidal Neurons. J. Neurosci. 29, 3233–3241.

42. Testa-Silva, G., Verhoog, M.B., Goriounova, N.B., Loebel, A., Hjorth, J., Baayen, J.C., De Kock, C.P.J., and Mansvelder, H.D. (2010). Human synapses show a wide temporal window for spike-timing-dependent plasticity. Front. Synaptic Neurosci. 2, 12.

43. Brzosko, Z., Schultz, W., and Paulsen, O. (2015). Retroactive modulation of spike timing-dependent plasticity by dopamine. Elife 4, e09685.

44. Bittner, K.C., Milstein, A.D., Grienberger, C., Romani, S., and Magee, J.C. (2017). Behavioral time scale synaptic plasticity underlies CA1 place fields. Science (80-.). 357, 1033–1036.

45. Turrigiano, G.G. (2017). The dialectic of Hebb and homeostasis. Philos. Trans. R. Soc. B Biol. Sci. 372, 20160258.

46. Stoppini, L., Buchs, P.A., and Muller, D. (1991). A simple method for organotypic cultures of nervous tissue. J. Neurosci. Methods 37, 173–82.

47. Gee, C.E., Ohmert, I., Wiegert, J.S., and Oertner, T.G. (2017). Preparation of Slice Cultures from Rodent Hippocampus. Cold Spring Harb. Protoc. 2017, 126–130.

48. Wiegert, J.S., Gee, C.E., and Oertner, T.G. (2017). Viral Vector-Based Transduction of Slice Cultures. Cold Spring Harb. Protoc. 2017, 131–134.

49. Wiegert, J.S., Gee, C.E., and Oertner, T.G. (2017). Single-Cell Electroporation of Neurons. Cold Spring Harb. Protoc. 2017, 135–138.

50. Suter, B.A., O’Connor, T., Iyer, V., Petreanu, L.T., Hooks, B.M., Kiritani, T., Svoboda, K., and Shepherd, G.M.G. (2010). Ephus: multipurpose data acquisition software for neuroscience experiments. Front. Neural Circuits 4, 100.

51. Schindelin, J., Arganda-Carreras, I., Frise, E., Kaynig, V., Longair, M., Pietzsch, T., Preibisch, S., Rueden, C., Saalfeld, S., Schmid, B., et al. (2012). Fiji: an open-source platform for biological-image analysis. Nat. Methods 9, 676–682.

